# The Development and Evaluation of a Fully Automated Markerless Motion Capture Workflow

**DOI:** 10.1101/2022.02.16.480655

**Authors:** Laurie Needham, Murray Evans, Logan Wade, Darren P. Cosker, Polly M. McGuigan, James L. Bilzon, Steffi L. Colyer

**Affiliations:** Centre for the Analysis of Motion, Entertainment Research and Applications, University of Bath, Bath, UK

**Keywords:** computer vision, pose estimation, validation, biomechanics, inverse kinematics, deep learning

## Abstract

This study presented a fully automated deep learning based markerless motion capture workflow and evaluated its performance against marker-based motion capture during overground running, walking and counter movement jumping. Multi-view high speed (200 Hz) image data were collected concurrently with marker-based motion capture (criterion data), permitting a direct comparison between methods. Lower limb kinematic data for 15 participants were computed using 2D pose estimation, our 3D fusion process and OpenSim based inverse kinematics modelling. Results demonstrated high levels of agreement for lower limb joint angles, with mean differences ranging between 0.1° - 10.5° for 6 DoF hip joint rotations, and 0.7° - 3.9° for knee and ankle rotations. These differences generally fall within the documented uncertainties of marker-based motion capture, suggesting that our markerless approach could be used for appropriate biomechanics applications. We used an open-source, modular and customisable workflow, allowing for integration with other popular biomechanics tools such as OpenSim. By developing open-source tools, we hope to facilitate the democratisation of markerless motion capture technology and encourage the transparent development of markerless methods. This presents exciting opportunities for biomechanics researchers and practitioners to capture large amounts of high quality, ecologically valid data both in the laboratory and in the wild.

## INTRODUCTION

The promise of markerless motion capture for biomechanics applications is by no means new (Colyer, Evans, Cosker, & Salo, 2018; Mündermann, Corazza, & Andriacchi, 2006). But despite being eagerly anticipated by the biomechanics community, markerless motion capture tools have never, until recently, provided a viable alternative to marker-based systems. Early generations of markerless motion capture typically made use of background subtraction techniques to extract silhouettes of the person of interest, to which a joint constrained, rigid body model could be fit via a mathematical minimisation routine (Corazza et al., 2006). Such an approach required a dense array of calibrated cameras and often required large amounts of human intervention to resolve errors in the model fitting process (Mündermann et al., 2006). However, thanks to the rapid maturation of deep-learning based pose estimation algorithms, a new generation of markerless motion capture technologies are emerging. Most notably, pose estimation using deep convolutional neural networks, which estimate a set of sparse key points in 2D images, represent an exciting emerging technology that has the potential to compliment and even replace current motion capture techniques (Kanko, Laende, Selbie, & Deluzio, 2021).

However, the naive application of 2D pose estimation methods may provide at best, planar estimates of joint centres and vector-based angles which are known to be error prone (Cronin, 2021; Kidzinski et al., 2020). Alternatively, when applied across multiple calibrated camera views, with a suitable 3D reconstruction and outlier removal algorithm, 2D pose estimation has been used to successfully detect and reconstruct the 3D position of human joint centres (Kanko, Laende, Davis, Selbie, & Deluzio, 2021; Nakano et al., 2020; Needham et al., 2021). When evaluated against marker-based motion capture data, differences of approximately 30-50 mm have been recorded for activities such as walking, running, jumping and throwing (Kanko et al., 2021; Nakano et al., 2020; Needham et al., 2021). These differences indicated that markerless approaches do not match the results of marker-based motion capture exactly, but arguably this should not be goal of markerless motion capture, as marker-based methods are not without their own biases and limitations e.g., soft tissue artefact (Miranda, Rainbow, Crisco, & Fleming, 2013) and marker placement reliability errors (Della Croce, Leardini, Chiari, & Cappozzo, 2005). Nevertheless, the agreement in joint centre location identification reported between markerless and marker-based systems generally falls within the known uncertainties for marker-based results when compared to gold standard biplanar videoradiography (Kessler et al., 2019; Miranda et al., 2013).

Determining accurate joint centre locations is fundamental to any analysis of human movement, regardless of the motion capture method being employed. However, most biomechanical analyses seek to move beyond planar descriptions, instead describing the position and orientation (pose) of a body segment with six degrees of freedom (6DoF). Raw data from any motion capture system is known to contain some degree of measurement artefact, for example, soft tissue artefact in marker data (Benoit, Damsgaard, & Andersen, 2015), keypoint jitter in pose estimation data (Needham, Evans, Cosker, & Colyer, 2021; Needham et al., 2021) or sensor drift in inertial measurement units (Rum et al., 2021). It is common practice to employ techniques such as inverse kinematics (IK), where a global optimisation process fits a rigid-body model with physically realistic joint constraints to noisy movement data (Lu & O’Connor, 1999). IK modelling has been shown to reduce the effects of measurement artefact (Aristidou & Lasenby, 2009; Begon, Andersen, & Dumas, 2018; Clement, Dumas, Hagemeister, & de Guise, 2015) in human movement data. A further advantage of IK modelling is the reduced requirement for three non-colinear points on a rigid body of interest to compute its pose. Such a feature is useful for pose estimation keypoint models which typically only detect joint centre locations (e.g., COCO and MPII keypoint models) and do not always include an additional off-axis point required for 6DoF segment pose to be directly inferred.

Currently, researchers are limited to closed-source, black-box and potentially expensive markerless motion capture tools. The democratisation of markerless technology will therefore require the development of a modular workflow using open-source computer vision tools including multi-camera calibrations, 2D pose estimation methods, 3D reconstruction methods, multi-person tracking and robust outlier detection, a suitable IK model and integration with other open-source biomechanics tools e.g., OpenSim (Delp et al., 2007). In this paper, we present an end-to-end automated workflow for computing 6DoF segment kinematics from calibrated multi-camera data. The aim of this study was to detail the development of our markerless motion capture system and rigorously evaluate it against marker-based motion capture during walking, running, and jumping.

## METHODS

### Markerless System Development

An overview of each stage of the markerless system’s workflow provided in Figure 1.

**Figure 1.**
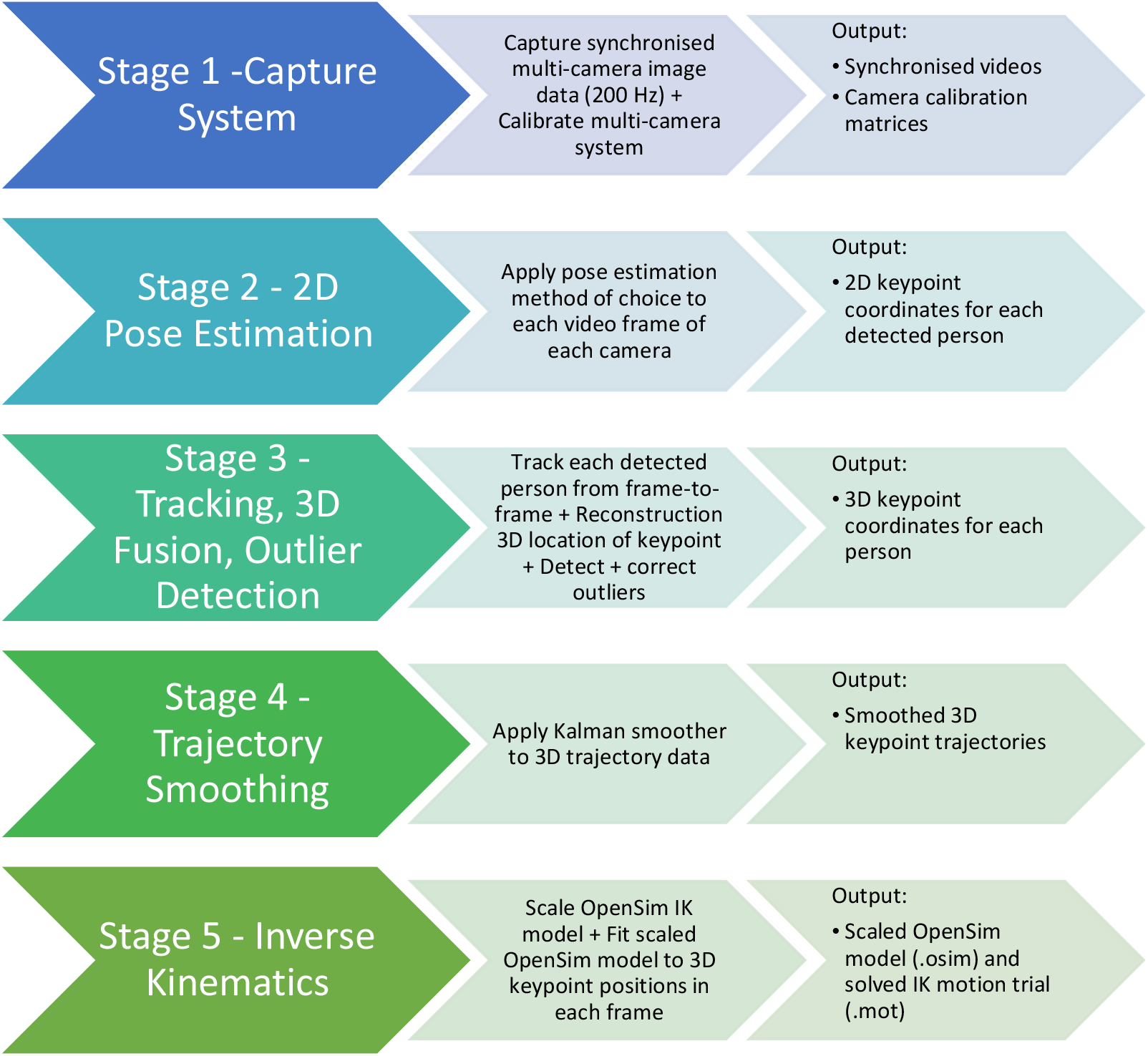
Flow chart depicting each stage our markerless motion capture workflow from recording data (stage 1) to producing a 3D IK constrained OpenSim model of the recorded motion (stage 5). The central column provides an overview of the processing required at each stage and the right-hand column provides details of the output from a given processing stage which becomes the input for the next stage. Each processing stage is described in detail in the methods section.

Stage 1 - Camera System – For this study we utilised nine TTL-pulse synchronised machine vision cameras (JAI sp5000c, JAI ltd, Denmark) capable of capturing HD images at 200 Hz. This provided the flexibility to create large capture volumes (∼10 × 10 × 3 m) with multiple views providing robustness against occlusions, however our approach can be used with as few as three cameras and more than nine cameras if computational and data storage resources are sufficient. Multi camera calibration was achieved using a binary circle pattern calibration board to initialise each camera’s intrinsic parameters and lens distortion coefficients (Zhang, 2000) The calibration board was then moved through the volume ensuring multiple shared observations between camera views. Sparse Bundle Adjustment (Triggs, McLauchlan, Hartley, & Fitzgibbon, 1999) was used to determine a set of globally optimum intrinsic and extrinsic camera parameters (Figure 2).

**Figure 2.**
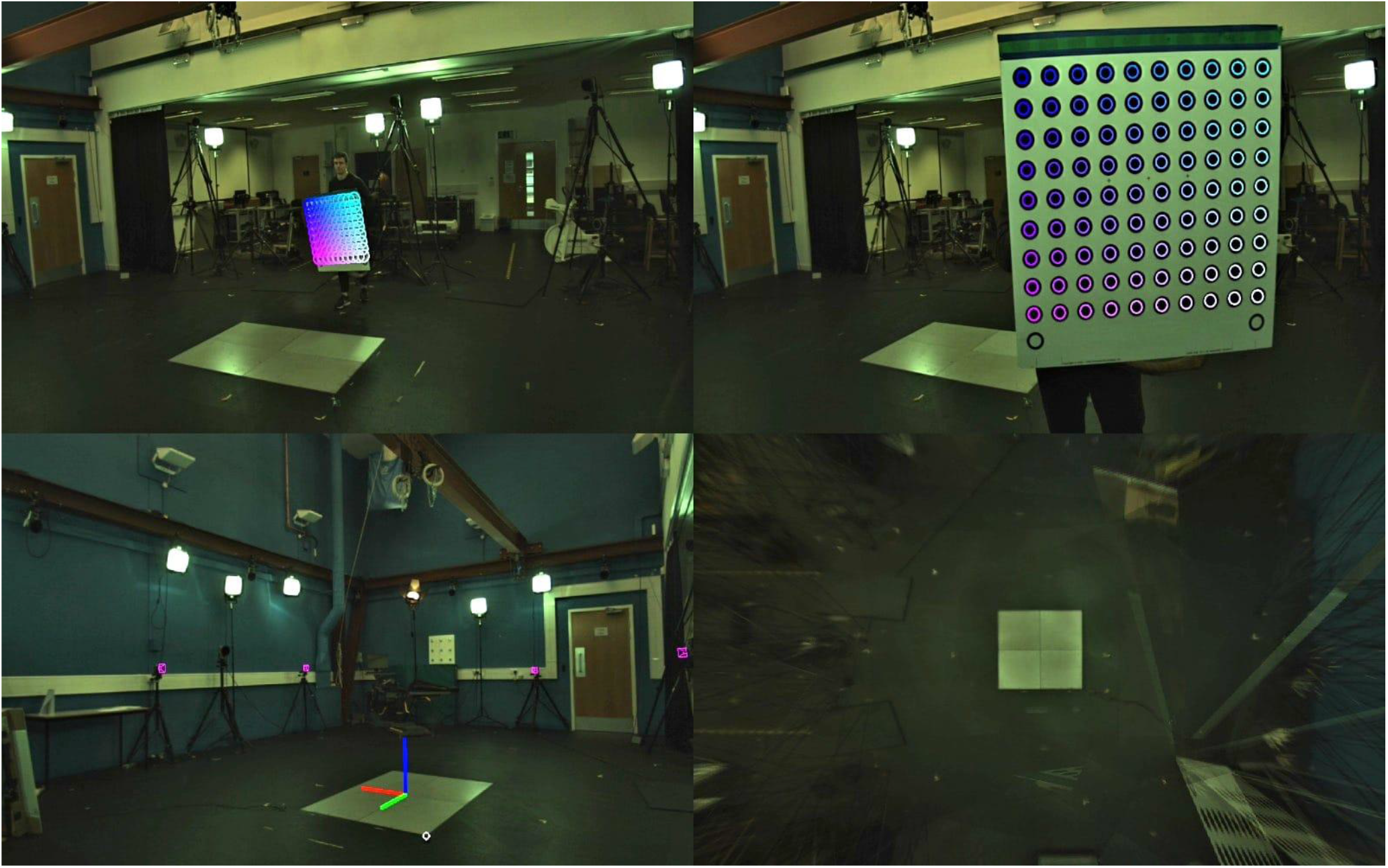
Example of the camera calibration procedure. In each camera view, where the calibration board’s grid of dots is detected, coloured circles are rendered onto the board (top-left, top right). Shared observations of these points, of known dimensions, allow for the estimation of each camera’s intrinsic parameters e.g., focal length, optical centre, and distortion coefficients, and extrinsic parameters e.g., each cameras position and orientation relative to the laboratory coordinate system. Lower-left shows calibrated camera locations (magenta boxes) and the global coordinate system location. As is seen in this figure, calibration quality can be verified by ensuring that the computed camera locations align with the actual camera locations as seen in each camera view. Lower right shows a synthetic overhead image where each camera view has been projected onto the ground plane. This image can be used to verify the ground plane calibration and, in this case, clearly shows the laboratory’s force plates in the centre of the plane.

Stage 2 - 2D Pose Estimation - 2D pose estimation of sparse keypoints (joint centres) in each camera view was achieved using an implementation of OpenPose (Cao, Simon, Wei, & Sheikh, 2017; Cao, Simon, Wei, Sheikh, & Ieee, 2017) and specifically, its ‘body_25’ model (Figure 3), as we have previously shown that this method performs well on biomechanics image data (Needham et al., 2021; Needham et al., 2021). However, the modular nature of our workflow means that most pose estimation methods could be utilised at this stage, facilitating new state-of-the-art network architectures, networks specialised to a specific task or object of interest, or networks trained on biomechanics specific datasets.

**Figure 3.**
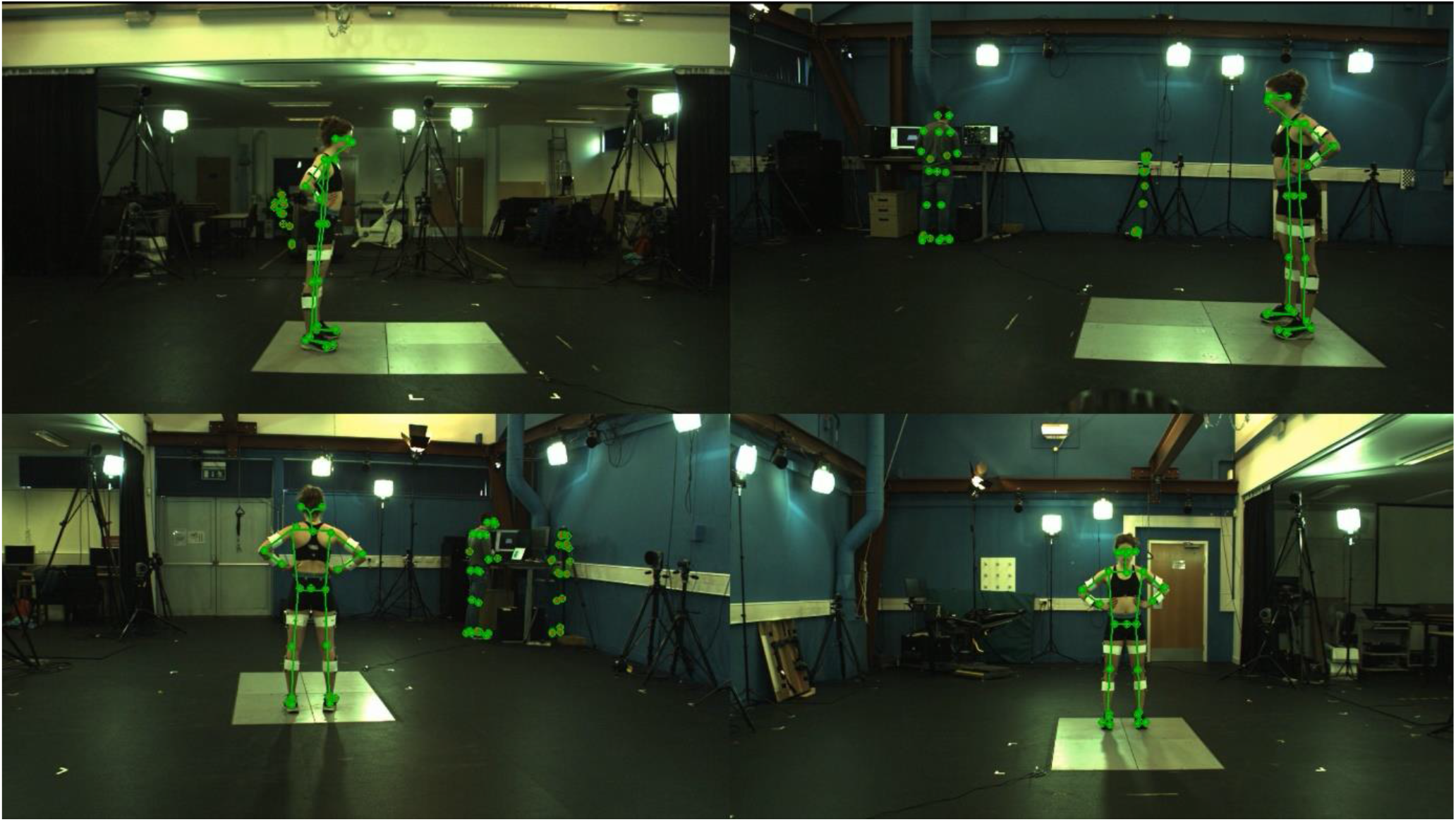
Example 2D joint centre detections and 3D joint centre reconstructions. 2D OpenPose detections for each camera view are rendered as green circles where the fill colour depicts the keypoint confidence values (e.g. red < 0.2, green > 0.2). Green lines depict 3D reconstructed joint centres following the tracking, fusion, outlier removal and Kalman smoothing stages. Examples of multi-person detections of people in the background and, of false-positive detections can be seen on tripods in the background. These detection challenges are efficiently and robustly handled by the tracking, 3D reconstruction and outlier detection stages.

Stage 3 - Tracking, 3D Fusion & Outlier Detection – Many pose estimation methods, including OpenPose, provide keypoint coordinates on the 2D image plane independently for each view at each time instance. To associate multi-person detections across views and track them through time, we used our previously developed approach based on occupancy maps which can efficiently associate multi-person detections between viewpoints and time instances (Needham et al., 2021). Once detections are associated between camera views, a 3D Euclidean space reconstruction of each person’s keypoints can be generated. Each 2D keypoint detection was back-projected using the appropriate camera calibration, accounting for non-linearities such as lens distortion, and resulting in a set of rays in the 3D space. The 3D reconstructed position of a given point was determined from the ‘optimal’, in a least squares sense, intersection of these rays (Slabaugh, Schafer, & Livingston, 2001). To handle common failures of the OpenPose detector such as misplaced limbs or contra-lateral label swaps, a robust RANSAC-like approach (Fischler & Bolles, 1981) was used to identify inlier and outlier rays for each keypoint, with the final reconstruction coming from only the inlier set assuming at least three inlier rays contributed to the reconstruction and the keypoint confidence value of each of these rays was greater than 0.3 (Figure 3).

Stage 4 - Trajectory Smoothing – A bi-directional Kalman filter which considered both previous and future states of the trajectory was used to provide a smooth output that was robust to outliers which bypassed the occupancy map and RANSAC processing stages. Previously, we have demonstrated this approach to be more effective for markerless pose estimation data than traditional low-pass filtering (Needham et al., 2021).

Stage 5 - Inverse Kinematics – 3D reconstructed keypoints were used to drive the motion of a constrained rigid-body kinematic model which could describe the position and orientation of each segment as a set of generalised coordinates. In this study, the marker-based and markerless models were scaled using a static calibration trial to negate scaling differences that could arise between the marker and markerless data and permit a more direct comparison (Figure 4). However, the markerless model could instead be scaled using median values taken during a movement trial, meaning that collecting a static calibration trial is not mandatory for this system. The model pose was globally optimised in each frame using OpenSim’s (Delp et al., 2007) IK tools accessed via the C++ API. Again, due to the modular nature of our workflow, other IK solvers could be implemented here. In this study the hip joints were given 3 DoF, the knee joints 1DoF (flexion/extension) and the ankle joint 2DoF (plantar dorsi/plantar flexion and ankle ad/abduction) which reflected the number of markerless key points that were available. Graphic Rendering – Details on how 3D joints centre locations (Figure 3) and 3D OpenSim models (Figure 5) were projected onto camera image planes for visualisation purposes are provided in the supplementary materials.

**Figure 4.**
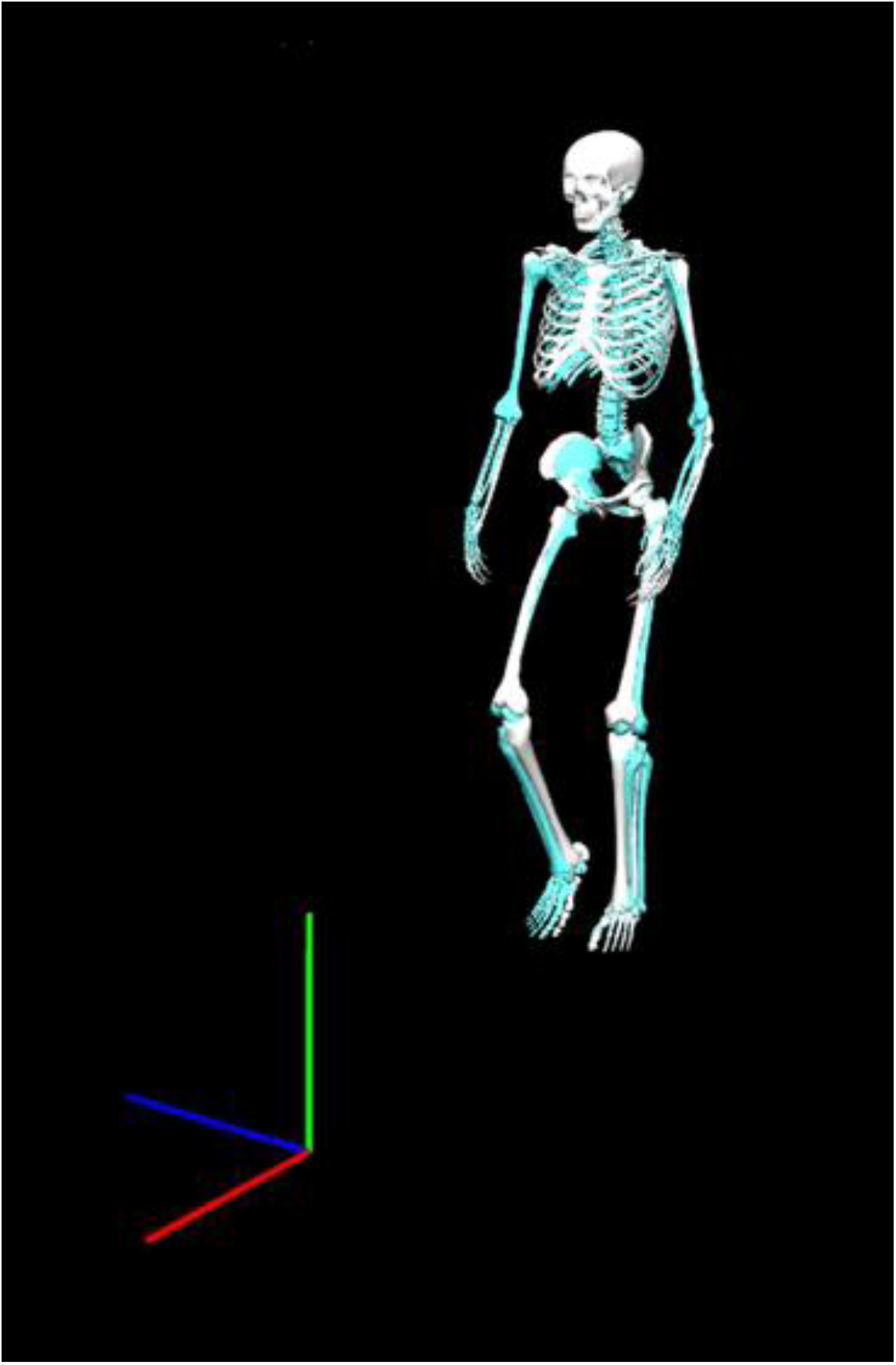
Example marker-based (white) and markerless (cyan) skeletal models fit to the same trial and video frame during walking.

**Figure 5.**
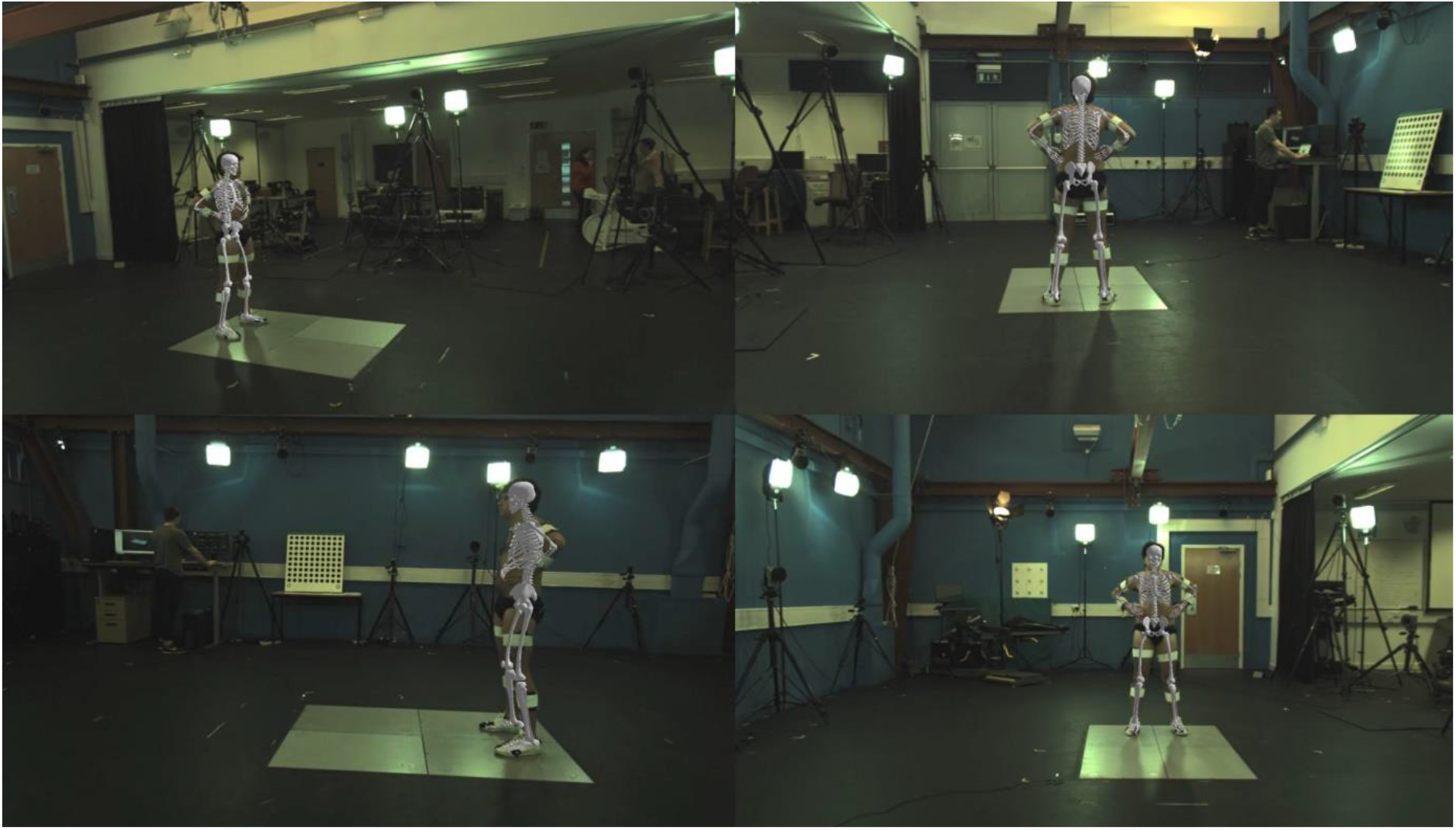
Example 3D IK skeletal model projected on select image planes and overlaid onto original image data. To verify the final markerless IK solution, the 3D OpenSim model has been projected into each camera view and overlaid onto the participant using the relevant camera calibration information.

**Figure 5.**
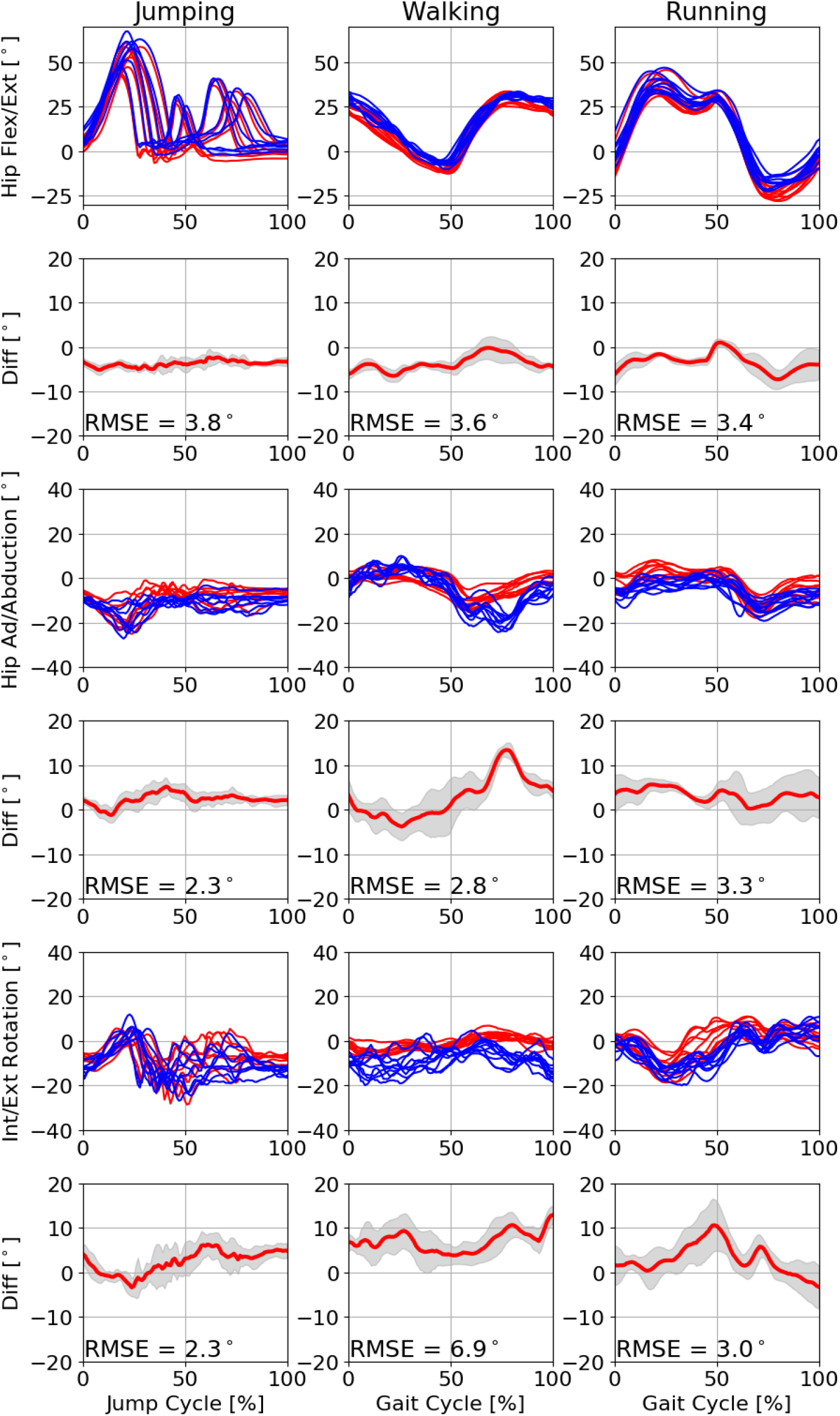
Rows 1, 3, 5 - Example time normalised marker-based (red) and markerless (blue) hip joint angles for a single participant (P06) during jumping (column 1), walking (column 2), and running (column 3). Rows 2, 4, 6 - Mean differences (red) ± SD (shaded area) for each corresponding joint angle figure with RMSE values embedded. RMSE values are given for the example participant.

### Criterion Data

15 participants (7 males, 8 females [1.71 ± 0.10 m, 73.7 ± 14.7 kg]) provided written informed consent. Each participant performed ten running trials at a self-selected speed, ten walking trials at self-selected speed and ten counter movement jumps with hands held on the waist. Motion data were captured concurrently using our markerless system and a criterion marker-based motion capture system.

Criterion data were captured using a 15-camera marker-based motion capture system (Oqus, Qualisys AB, Gothenburg, Sweden). Motion capture systems were time-synchronised by means of a periodic TTL-pulse generated by the custom system’s master frame grabber to achieve a sampling frequency of 200 Hz in both systems. This ensured that frames were captured by all cameras in unison without drift. A right-handed coordinate system was defined for both systems by placing a Qualisys L-Frame in the centre of the capture volume. To refine the alignment of each system’s Euclidean space, a single marker was moved randomly through the capture volume and tracked by both systems. These tracked points provided two sets of corresponding point clouds with which the spatial alignment could be optimised in a least-squares sense. To capture criterion data, a full body marker set comprising 44 individual markers and four clusters were attached to each participant to create a full body 6DoF model (bilateral feet, shanks and thighs, pelvis and thorax, upper and lower arms, and hands) along with rigid clusters placed bilaterally on the upper arm and forearms and upper and lower legs. Following labelling and gap filling of trajectories (Qualisys Track Manager v2019.3, Qualisys, Gothenburg, Sweden), marker-based data were filtered (Butterworth low-pass, cut-off 12 Hz) and used to scale and drive the motion of an OpenSim IK model. Walking and running data were time normalised to a gait cycle (touch-down to the proceeding touch-down of the same foot) where touch-down and toe-off events were determined using a 20 N vertical force threshold. Jumping data were time normalised from first movement (first instance of the net vertical force dropping and staying below zero for 50 frames) to quiet standing (net vertical force returned to and stayed within 3x SDs of zero after landing).

### System Evaluation

To evaluate agreement between marker and markerless systems, hip (flexion/extension, ad/abduction, int/external rotation), knee (flexion/extension), and ankle (plantar/dorsiflex) Euler angles from both systems were compared using root mean squared errors (RMSE), Bland-Altman analyses (Bland & Altman, 1999) and linear fit models (Iosa et al., 2014).

## RESULTS

An example result for the markerless system is shown in Figure 4 (marker vs markerless skeletal OpenSim models) and Figure 5 (OpenSim model overlaid onto original images) for context of relative differences. Good levels of agreement can be observed between marker-based and markerless results although clearly, there is not perfect agreement between systems.

For hip joint angles, good agreement was observed between both systems, particularly for hip flexion/extension and hip ad/abduction (Figure 5 – single participant example, Table 1 – group level statistic) with Bland-Altman analysis revealing low bias (< 3°) and low random error (< 3°) during all activities. During all activities, the largest differences between marker-based and markerless hip joint angles were observed for internal/external rotation, most notably during walking with mean differences of 10.5 ± 2.9 ° (Table 1). However, the low random error observed suggests that a portion of these differences were systematic in nature.

**Table 1:** Bland-Altman analysis and linear fit model results for differences between marker-based and markerless hip angles for jumping, walking and running activities.

| Joint Angle | Activity | Mean Difference<br>(Bias) (°) | ± SD | 95% LoA | R<sup>2</sup> |
| --- | --- | --- | --- | --- | --- |
| Hip Flexion/Extension | Jumping | -2.8 | 2.2 | -7.1 - 1.5 | 0.72 |
|  | Walking | -0.1 | 2.6 | -5.1 - 5.1 | 0.13 |
|  | Running | -2.5 | 2.3 | -7.1 - 2.1 | 0.59 |
| Hip Ad/Abduction | Jumping | 1.7 | 2.1 | -2.3 - 5.7 | 0.61 |
|  | Walking | 3.7 | 2.5 | -1.2 - 8.6 | 0.11 |
|  | Running | 4.1 | 1.8 | 0.6 - 7.7 | 0.29 |
| Hip Internal/External<br>Rotation | Jumping | 5.6 | 3.5 | -1.3 - 12.5 | 0.34 |
|  | Walking | 10.5 | 2.9 | 4.8 - 16.2 | 0.61 |
|  | Running | 6.8 | 3.7 | -0.6 - 14.14 | 0.27 |

The knee joint exhibited good agreement between marker-based and markerless systems (Figure 6, Table 2) with Bland-Altman analysis revealing low bias (< 3°) and low random error (< 3°) during all activities. Despite low bias and random error values at a group level (Table 2) the largest joint angle differences at an individual level were observed for ankle dorsi/plantar-flexion during peak dorsiflexion for the participant example given in Figure 6, during walking. Here the marker-based system consistently estimated lower magnitudes of dorsi-flexion when compared to the markerless results.

**Figure 6.**
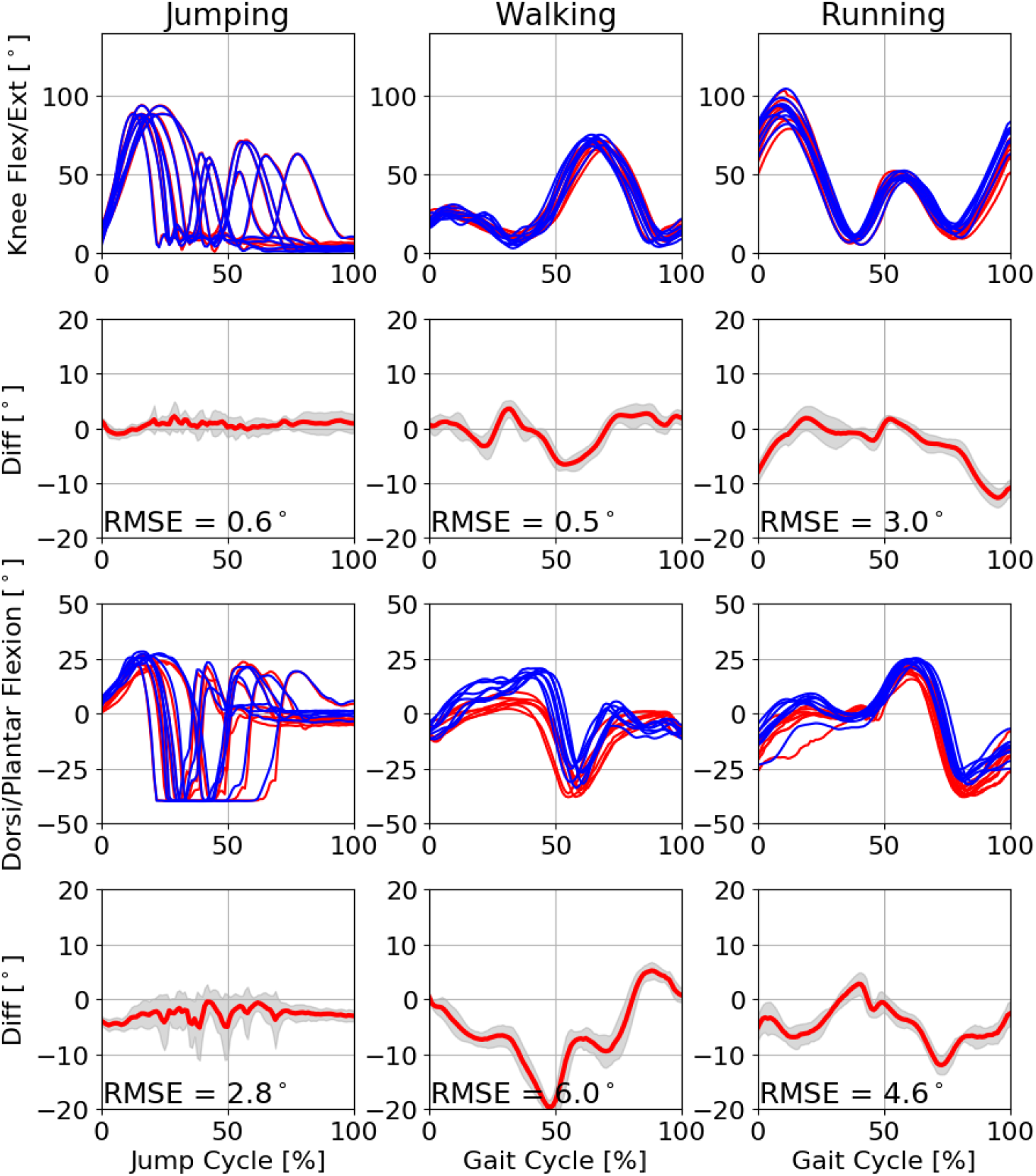
Rows 1, 3, 5 - Example time normalised marker-based (red) and markerless (blue) knee and ankle joint angles for a single participant (P06) during jumping (column 1), walking (column 2), and running (column 3). Rows 2, 4, 6 - Mean differences (red) ± SD (shaded area) for each corresponding joint angle figure with RMSE values embedded. RMSE values are given for the example participant.

**Table 2:**
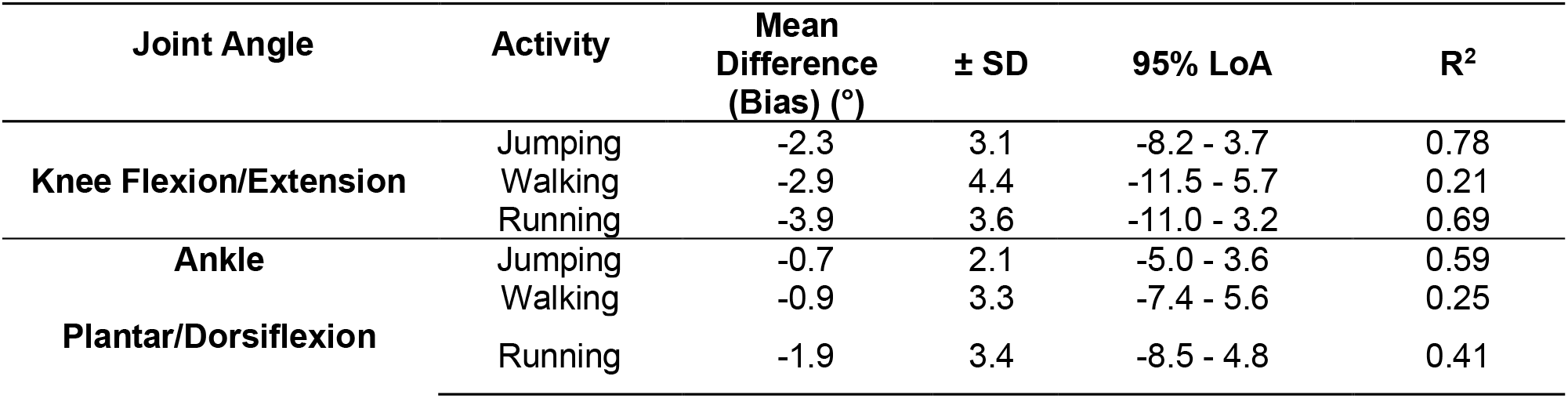
Bland-Altman analysis results for differences between marker-based and markerless knee and ankle angles.

## DISCUSSION

This study presents an end-to-end workflow for markerless motion capture and provides an evaluation against a marker-based system, focusing on lower limb joint angles during counter-movement jumping, overground walking and overground running. Our markerless motion capture system was able to capture kinematics within the known uncertainties of marker-based motion capture, most notably for sagittal and frontal plane motion (Table 1 and 2).

Hip flexion/extension consistently demonstrated high levels of agreement (mean differences < 3° - Table 1) between marker and markerless systems. Larger differences were observed during running and walking for hip ad/abduction (mean differences ∼ 4° - Table 1) and internal/external rotation (mean differences up to 10° - Table 1). The high levels of agreement for hip flexion/extension demonstrate that our markerless approach can achieve excellent segment pose estimation. This is perhaps due to the presence of favourable markerless keypoints for this axis of rotation (i.e. hip and knee joint centres) and the large range of motion reducing the requirement for precision in both systems. Where larger mean differences were observed for hip rotations, offsets between systems can be attributed to systematic differences that have previously been reported for joint centre locations (Needham et al., 2021). Additionally, in the markerless approach, there was a greater reliance on IK modelling to determine internal/external rotation which, due to an absence of an additional off-axis point on the thigh segment, is determined entirely by the pose of segments distal to the knee joint. Thus when the knee was in a more flexed position, we observed better agreement for internal/external rotations of the hip, which aligns with the findings of other IK studies (Begon et al., 2018; Kainz et al., 2017; Mantovani & Lamontagne, 2017). Future pose estimation models might address this issue by placing additional off-axis points on either anatomically meaningful landmarks such as the medial and lateral knee, and/or less anatomically meaningful points such as the mid-anterior and mid-posterior surface of the thigh segment. It is important to note however, that an unknown proportion of the differences reported for the hip joint rotations are likely due to errors arising from marker-based soft tissue artefact which itself has been attributed to errors of up to 12° (D’Isidoro, Brockmann, & Ferguson, 2020; Fiorentino, Atkins, Kutschke, Foreman, & Anderson, 2020), particularly during high-frequency impacts such as jumping and running touchdown events (Miranda et al., 2013).

High levels of agreement were also observed for the knee joint, with mean differences less than 4° (Table 2), again demonstrating that our markerless approach can produce results aligning with marker-based systems, and commercial markerless systems (Kanko et al., 2021). Larger differences were observed in knee flexion/extension for running with some peak values greater than 10° occurring around touchdown and early stance. These differences may be due in part to systematic differences in joint centre locations computed at the 2D pose estimation stage, resulting in different knee joint centre locations of up to 40 mm (Needham et al., 2021). However, it is also likely that an unknown proportion of the differences observed were due to error in the marker-based data, with soft tissue artefact contributing up to 7° of error during touchdown and early stance (Miranda et al., 2013). The IK knee model used in this study was constrained to a single axis of rotation, however, it is possible to permit additional rotations such as knee ad/abduction. However, the current OpenPose keypoint model could not provide enough information pertaining to the pose of the thigh and shank segments to permit accurate and reliable knee rotations beyond the sagittal plane.

Differences exhibited for ankle dorsi/plantar flexion angles were likely the result of several factors. Firstly, sub-optimal pose estimation at the 2D detection stage when locating distal foot keypoints in each images. Indeed, the poorest keypoint detections for OpenPose were typically observed for the toe and heel points (Needham et al., 2021) and were likely caused by these keypoints being trained on a much smaller subsample of the COCO dataset (approximately 14,000 images (Cao, Simon, Wei, & Sheikh, 2017)). Secondly, systematic differences in knee and hip joint centre locations may have propagated down the IK chain, resulting in poorer pose estimation of the most distal segments. Such errors are a product of IK’s global optimisation, meaning that errors in one joint can affect all joints (Kainz et al., 2017). However, as the same IK optimisations were used for both marker and marker-less data in this study, it is possible that a proportion of the differences observed are due to errors in both systems.

The results of this study are highly promising and the differences between systems typically fell within the known uncertainties of marker-based motion capture. For example, errors of up to ∼9° have been reported for marker-based motion capture when compared to biplanar videoradiography for knee flexion/extension (Miranda et al., 2013) and ankle joint rotations (Kessler et al., 2019). As such our system could be employed for applications where the present accuracy meets the required minimum detectable change for a given research application. Marker-based motion capture represents the current de-facto standard in many biomechanics applications and its limitations including soft tissue artefact, marker placement reliability, and joint axis crosstalk have been well documented (Begon et al., 2018; Benoit et al., 2015; Kainz et al., 2017; Kessler et al., 2019; Miranda et al., 2013). It is important that the same rigorous evaluation processes be applied to markerless motion capture so that researchers and practitioners can make informed decisions about their strengths and weaknesses.

To facilitate improved pose estimation performance for biomechanics applications, there is ultimately a need for large open-access image datasets which are annotated with high anatomical accuracy, reliability, and objectivity (Cronin, 2021). Ensuring that each body segment of interest is annotated with at least three non-colinear points, ideally more, will facilitate the estimation of segment poses with greater accuracy, robustness, and additional degrees of freedom. Furthermore, such datasets would facilitate reduced reliance on global pose optimisation methods (e.g., IK solvers) and allow independent segment pose optimisation solutions to be implemented, giving users greater flexibility.

This work takes a step towards democratising markerless motion capture by developing an open-source, modular, and transparent set of tools. Open-source biomechanics tools such as OpenSim (Delp et al., 2007), Pyomeca (Martinez, Michaud, & Begon, 2020) and Biomechanical ToolKit (Barre & Armand, 2014) have had made a widespread positive impact upon the biomechanics community and the nature of open-source software presents several advantages over proprietary systems. Cost limitations are reduced, thus increasing accessibility, and researchers can access the source code directly to determine how outputs have been computed. The modular nature of our system provides the flexibility for users to customise each stage to their specific needs, for example, implementing custom trained 2D pose estimation models or modified IK models. Integrating pose outputs directly into other open-source biomechanics tools, such as OpenSim, may also facilitate a range of simulation-based studies in the future. In due course, as markerless motion capture technologies continue to develop, we should ask ourselves if benchmarking against marker-based technologies remains the best option or whether we should seek to establish new methods of acquiring high fidelity ground-truth information that can be used to advance measurement technologies and most importantly allow us to solve current problems and answer new research questions.

## CONCLUSION

We have presented a novel, fully automated and open-source markerless motion capture workflow. Evaluation results were comparable to marker-based motion capture and commercial markerless systems. A need for large scale, open access biomechanics pose datasets was identified alongside the need for better evaluation datasets which will facilitate the development of the next generation of markerless motion capture systems. Markerless motion capture technologies are maturing at a rapid pace and present a promising approach to collecting biomechanics data in the wild.

## ACKNOWLEDGEMENT

This research was funded by CAMERA, the RCUK Centre for the Analysis of Motion, Entertainment Research and Applications, EP/M023281/1 and EP/T014865/1. Thank you to Dr Nicos Haralabidis and James Cowburn for their invaluable OpenSim knowledge which helped us hack our way through the OpenSim C++ API with relative ease.

