## Appendix 1 for "The Development and Evaluation of a Fully Automated Markerless Motion Capture Workflow"

Graphic Rendering Process – To qualitatively inspect and visualise the 3D markerless results for each frame, 3D joint centre locations were projected onto camera image planes using the relevant camera calibration information. This allowed for the 3D joint centre locations to be overlaid onto the video frames. To overlay the markerless IK skeletal model results onto video frames, we iteratively extracted the pose and scaling information of the OpenSim model's bodies (segments), composing the pose parameters (Euler angles and offset translations) into homogeneous 4x4 transformation matrices. Each model segment is represented by a 3D mesh and loaded from .vtp files (Visualization Toolkit) provided with OpenSim. A custom OpenGL renderer which uses projections derived from the camera calibrations was then used to render the posed body segments on top of the corresponding video frame, allowing the final IK constrained 3D solution to be visualised (Figure 5).
